# Genomic analysis of family UBA6911 (Group 18 Acidobacteria) expands the metabolic capacities of the phylum and highlights adaptations to terrestrial habitats

**DOI:** 10.1101/2021.04.09.439258

**Authors:** Archana Yadav, Jenna C. Borrelli, Mostafa S. Elshahed, Noha H. Youssef

## Abstract

Approaches for recovering and analyzing genomes belonging to novel, hitherto unexplored bacterial lineages have provided invaluable insights into the metabolic capabilities and ecological roles of yet-uncultured taxa. The phylum Acidobacteria is one of the most prevalent and ecologically successful lineages on earth yet, currently, multiple lineages within this phylum remain unexplored. Here, we utilize genomes recovered from Zodletone spring, an anaerobic sulfide and sulfur-rich spring in southwestern Oklahoma, as well as from multiple disparate soil and non-soil habitats, to examine the metabolic capabilities and ecological role of members of the family UBA6911 (group18) Acidobacteria. The analyzed genomes clustered into five distinct genera, with genera Gp18_AA60 and QHZH01 recovered from soils, genus Ga0209509 from anaerobic digestors, and genera Ga0212092 and UBA6911 from freshwater habitats. All genomes analyzed suggested that members of Acidobacteria group 18 are metabolically versatile heterotrophs capable of utilizing a wide range of proteins, amino acids, and sugars as carbon sources, possess respiratory and fermentative capacities, and display few auxotrophies. Soil-dwelling genera were characterized by larger genome sizes, higher number of CRISPR loci, an expanded carbohydrate active enzyme (CAZyme) machinery enabling de-branching of specific sugars from polymers, possession of a C1 (methanol and methylamine) degradation machinery, and a sole dependence on aerobic respiration. In contrast, non-soil genomes encoded a more versatile respiratory capacity for oxygen, nitrite, sulfate, trimethylamine N-oxide (TMAO) respiration, as well as the potential for utilizing the Wood Ljungdahl (WL) pathway as an electron sink during heterotrophic growth. Our results not only expand our knowledge of the metabolism of a yet-uncultured bacterial lineage, but also provide interesting clues on how terrestrialization and niche adaptation drives metabolic specialization within the Acidobacteria.

**Importance:** Members of the Acidobacteria are important players in global biogeochemical cycles, especially in soils. A wide range of Acidobacterial lineages remain currently unexplored. We present a detailed genomic characterization of genomes belonging to the family UBA6911 (also known as group 18) within the phylum Acidobacteria. The genomes belong to different genera and were obtained from soil (genera Gp18_AA60 and QHZH01), freshwater habitats (genera Ga0212092 and UBA6911), and anaerobic digestor (Genus Ga0209509). While all members of the family shared common metabolic features, e.g. heterotrophic respiratory abilities, broad substrate utilization capacities, and few auxotrophies; distinct differences between soil and non-soil genera were observed. Soil genera were characterized by expanded genomes, higher numbers of CRISPR loci, larger carbohydrate active enzyme (CAZyme) repertoire enabling monomer extractions from polymer side chains, and methylotrophic (methanol and methylamine) degradation capacities. In contrast, non-soil genera encoded more versatile respiratory capacities for utilizing nitrite, sulfate, TMAO, and the WL pathway, in addition to oxygen as electron acceptors. Our results not only broaden our understanding of the metabolic capacities within the Acidobacteria, but also, provide interesting clues on how terrestrialization shaped Acidobacteria evolution and niche adaptation.

## Introduction

Our appreciation of the scope of phylogenetic and metabolic diversities within the microbial world is rapidly expanding. Approaches enabling direct recovery of genomes from environmental samples without the need for cultivation allow for deciphering the metabolic capacities and putative physiological preferences of yet-uncultured taxa (1–9). Further, the development of a genome-based taxonomic framework that incorporates environmentally-sourced genomes (10) has opened the door for phylocentric (lineage-specific) studies. In such investigations, comparative analysis of genomes belonging to a target lineage is conducted to determine its common defining metabolic traits, the adaptive strategies of its members to various environments, and evolutionary trajectories of and patterns of gene gain/loss across this lineage.

Members of the phylum Acidobacteria are one of the most dominant, diverse and ecologically successful lineages within the bacterial domain (11–16). Originally proposed to accommodate an eclectic group of acidophiles (17), aromatic compound degraders and homoacetogens (18), and iron-reducers (19), it was subsequently identified as a soil-dwelling bacterial lineage in early 16S rRNA gene-based diversity surveys (20–22). Subsequent 16S rRNA studies have clearly shown its near-universal prevalence in a wide range of soils, where it represents 5-50% of the overall community (23, 24).

Various taxonomic outlines have been proposed for the Acidobacteria. Genome-based classification by the Genome Taxonomy Database (GTDB, (r95, October 2020) (10) splits the phylum into 14 classes, 34 orders, 58 families, and 175 genera. This classification broadly, but not always, corresponds to 16S rRNA gene-based taxonomic schemes in SILVA (25), the 26 groups (subdivisions) classification scheme (26), and the most recently proposed refined class/order classification scheme (27) (Table S1). Regardless, a strong concordance between habitat and phylogeny was observed within most lineages in the Acidobacteria. Some lineages, e.g. groups 1, 3 (both in Class Acidobacteriae in GTDB), and group 6 (Class Vicinamibacteria in GTDB), have predominantly been encountered in soils, while others, e.g. groups 4 (Class Blastocatellia in GTDB), 8 (Class Holophagae in GTDB), and 23 (Class Thermoanaerobaculia in GTDB), are more prevalent in non-soil habitats (11).

Genomic analysis of cultured (11) and uncultured metagenome-assembled genomes (MAGs) and single cell genomes (SAGs) (28–30) representatives of the phylum Acidobacteria has provided valuable insights into their metabolic capacities and lifestyle. However, the majority of genomic (and other -omics) approaches have mostly focused on cultured, and yet-uncultured genomes of soil Acidobacteria (31–33). Genomic-based investigations of non-soil Acidobacteria pure cultures (34–37), or MAGs (38, 39) are more limited and, consequently, multiple lineages within the Acidobacteria remain unexplored.

We posit that genomic analysis of hitherto unexplored lineages of Acidobacteria would not only expand our knowledge of their metabolic capacities, but also enable comparative genomic investigation on how terrestrialization and niche adaptation shaped the evolutionary trajectory and metabolic specialization within the phylum. To this end, we focus on a yet-uncultured lineage in the Acidobacteria: Family UBA6911 (Subdivision 18 in (21), Class 1-2 in (27), and group 18 in SILVA database release 138.1 (25)). We combine the analysis of genomes recovered from Zodletone spring, an anaerobic sulfide and sulfur-rich spring in southwestern Oklahoma, with available genomes from multiple disparate soil and non-soil habitats. Our goal was to: understand the metabolic capacities, physiological preferences, and ecological role of this yet-uncharacterized group, and utilize the observed genus-level niche diversification to identify genomic changes associated with the terrestrialization process.

## Materials and Methods

### Sample collection, DNA extraction, and metagenomic sequencing

Samples were collected from the anaerobic source sediments in Zodletone Spring, a sulfide and sulfur rich spring in western Oklahoma’s Anadarko Basin (N34.99562° W98.68895°). The ecology, geochemistry, and phylogenetic diversity of the various locations within the spring has previously been described (40–43). The sediments were collected into sterile 50 mL polypropylene plastic tubes using sterile spatulas, transferred to the laboratory (within 2 hours) on ice and immediately processed for DNA extraction using the DNeasy PowerSoil kit (Qiagen, Valencia, CA, USA) according to manufacturer protocols. DNA was sequenced on the Illumina HiSeq 2500 platform using the services of a commercial provider (Novogene, Beijing, China). Sequencing produced 281.0 Gbp of raw data that were assembled using MegaHit (44), and binned using both Metabat (45) and MaxBin2 (46), followed by selection of the highest quality bins using DasTool (47). GTDB-Tk (48) (v 1.3.0) was used for the taxonomic classification of the bins using the classification workflow option -classify_wf, and 4 bins belonging to Acidobacteria Family UBA6911 were selected for further analysis. In addition, 18 genomes belonging to Family UBA6911 were selected from the recently released 52,515 genomes in the earth microbiome catalogue collection (49) available through the IMG/M database. These genomes were binned from peatland soils in Minnesota USA (4 genomes) (50, 51), an anaerobic biogas reactor in Washington, USA (11 genomes) (52), Cone Pool hot spring microbial mat in California, USA (2 genomes), and White Oak River estuary sediment in North Carolina, USA (1 genome) (53). Five other genomes belonging to Family UBA6911 were available from GenBank and were obtained in previous studies. These included 4 genomes binned from the Angelo Coastal Range Reserve in Northern California (30), and 1 genome binned from Noosa river sediments (Queensland, Australia) (54).

### Genomes quality assessment and general genomic features

Genome completeness, and contamination were assessed using CheckM (v 1.0.13) (55). All genomes included in this study were of medium or high quality with >70% completion and <10% contamination (Table S2). Designation as medium or high-quality drafts was based on the criteria set forth by MIMAGs (56). The 5S, 16S, and 23S rRNA sequences were identified using Barrnap 0.9 (https://github.com/tseemann/barrnap). tRNA sequences were identified and enumerated with tRNAscan-SE (v 2.0.6, May 2020) (57). Genomes were mined for CRISPR and Cas proteins using the CRISPR/CasFinder (58).

### Phylogenomic analysis

Taxonomic classification using the GTDB taxonomic framework was conducted using the classification workflow option -classify_wf within the GTDB-Tk (48). Phylogenomic analysis was conducted using the 120 single-copy marker genes (10) concatenated alignment that is generated by GTDB-Tk (48). Maximum-likelihood phylogenetic trees were constructed in RAxML (v 8.2.8) (59) with the PROTGAMMABLOSUM62 model and default settings. Representatives of all other Acidobacteria classes were included in the analysis (Figure 1); and *Chloroflexus aggregans* (GCF_000021945.1) was used as the outgroup. As expected, all 28 genomes were classified to the Family UBA6911 within the UBA6911 class of the Acidobacteria, but only 14 genomes were classified by the GTDB to the genus level into 2 genera gp18_AA60, and UBA6911, while the remaining genomes were unclassified at the genus level. To further assign these genomes to putative genera, average amino acid identity (AAI), and shared gene content (SGC) were calculated using the AAI calculator [http://enve-omics.ce.gatech.edu/]). Based on these values, we propose assigning these 14 unassigned genomes were assigned into three putative novel genera. We propose the names QHZH01 (1 genome from grassland soil), Ga0212092 (2 genomes from Cone pool microbial mat), and Ga0209509 (10 genomes from an anaerobic gas digestor (WA, USA), and 1 Zodletone sediment genome) based on the assembly accession number of the most complete genome within each genus (Tables 1 and S2).

**Figure 1.**
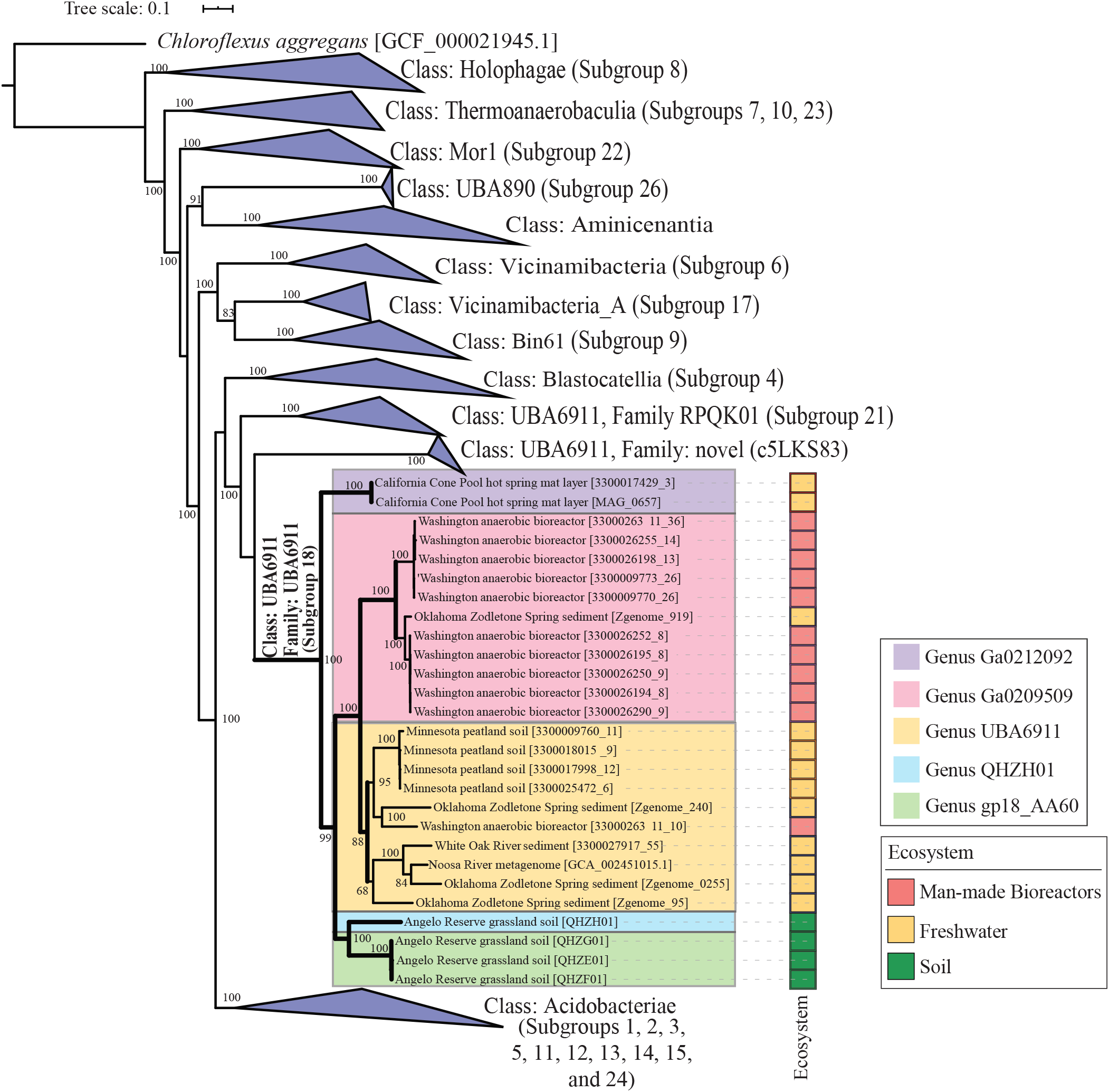
Maximum likelihood phylogenetic tree based on concatenated alignment of 120 single-copy genes from all Acidobacteria classes in GTDB r95. Family UBA6911, (Acidobacteria subgroup 18), is shown unwedged with thick branches. Branches are named by the Bin name between brackets (as shown in Table 1) and with the ecosystem from which they were binned. The five genera described here are color coded as shown in the legend. Bootstrap values (from 100 bootstraps) are displayed for branches with ≥70% support. The tree was rooted using *Chloroflexus aggregans* (GCF_000021945.1) as the outgroup. The tracks to the right of the tree represent the ecosystem from where the genomes were binned (color coded as shown in the legend). All other Acidobacteria classes are shown as wedges with the corresponding subgroup number(s) between parentheses.

**Table 1.** Binning sources, similarity statistics, and general genomics features for the genomes analyzed in this study.

| Genus name | Bin name | Binning Source | Similarity statistics |  | Binned genome size | Estimated Genome size | % protein coding bases | % GC content | No. of CRISPRs | Average gene length (bp) | Total number of Genes | Total number of protein-coding genes | Accession number | References |
| --- | --- | --- | --- | --- | --- | --- | --- | --- | --- | --- | --- | --- | --- | --- |
|  |  |  | AAI (average ± SD) | SGC (average ± SD) |  |  |  |  |  |  |  |  |  |  |
| Genus Ga0212092 | 3300022548-MAG_0657 | Cone Pool hot spring mat layer | 54.09 ± 0.98 | 40.34 ± 5.3 | 3.74 Mbp | 4.33 Mbp | 90.08% | 65.68% | 5 | 1021 | 3349 | 3303 | Ga0212092<sup>a</sup> | Unpublished* |
|  | 3300017429_3 |  |  |  | 3.82 Mbp | 4.56 Mbp | 90.05% | 65.60% | 4 | 1018 | 3424 | 3376 | Ga0185343<sup>a</sup> | Unpublished* |
| Genus QHZH01 | QHZH01 | Angelo reserve soil | 54.63 ± 5.62 | 35.97 ± 5.12 | 6.00 Mbp | 8.01 Mbp | 90.41% | 66.73% | 2 | 895 | 6088 | 6058 | QHZH01<sup>b</sup> | 30 |
| Genus Gp18_AA60 | QHZE01 | Angelo reserve soil | 55.32 ± 5.08 | 37.1 ± 4.46 | 7.54 Mbp | 8.12 Mbp | 84.85% | 58.70% | 18 | 985 | 6542 | 6497 | QHZE01<sup>b</sup> | 30 |
|  | QHZF01 |  |  |  | 7.06 Mbp | 8.42 Mbp | 84.49% | 58.85% | 25 | 916 | 6546 | 6508 | QHZF01<sup>b</sup> | 30 |
|  | QHZG01 |  |  |  | 7.33 Mbp | 7.65 Mbp | 84.45% | 58.87% | 31 | 980 | 6360 | 6314 | QHZG01<sup>b</sup> | 30 |
| Genus UBA6911 | 3300026311_10 | Anaerobic biogas reactor | 53.59 ± 4.66 | 38.83 ± 7.76 | 4.38 Mbp | 4.71 Mbp | 92.16% | 56.50% | 3 | 1002 | 4076 | 4029 | Ga0209723<sup>a</sup> | 52 |
|  | 3300009760_11 | Peatland |  |  | 3.10 Mbp | 5.17 Mbp | 84.98% | 52.10% | 0 | 827.4 | 3218 | 3184 | Ga0116131<sup>a</sup> | 50, 51 |
|  | 3300017998_12 | Peatland |  |  | 4.18 Mbp | 5.15 Mbp | 87.18% | 52.70% | 4 | 866 | 4262 | 4211 | Ga0187870<sup>a</sup> | 50, 51 |
|  | 3300025472_6 | Peatland |  |  | 4.15 Mbp | 5.61 Mbp | 86.20% | 52.60% | 4 | 838 | 4309 | 4267 | Ga0208692<sup>a</sup> | 50, 51 |
|  | 3300018015_9 | Peatland |  |  | 4.69 Mbp | 5.91 Mbp | 87.26% | 52.80% | 2 | 831 | 4956 | 4923 | Ga0187866<sup>a</sup> | 50, 51 |
|  | 3300027917_55 | White Oak River sediment |  |  | 3.87 Mbp | 4.43 Mbp | 84.76% | 48.90% | 9 | 898 | 3695 | 3656 | Ga0209536<sup>a</sup> | 53 |
|  | GCA_002451015.1 | Noosa river |  |  | 4.94 Mbp | 5.72 Mbp | 85.35% | 51.60% | 1 | 911 | 4657 | 4632 | DKBB01<sup>b</sup> | 54 |
|  | Zgenome_0255 | Zodlton Spring sediment |  |  | 3.75 Mbp | 5.12 Mbp | 87.68% | 57.60% | 19 | 986.9 | 3252 | 3220 | JAFGAO01<sup>b</sup> | This study |
|  | Zgenome_240 |  |  |  | 6.51 Mbp | 6.81 Mbp | 87.10% | 52.30% | 5 | 1010.5 | 5666 | 5615 | JAFGIY01<sup>b</sup> | This study |
|  | Zgenome_95 |  |  |  | 5.45 Mbp | 5.85 Mbp | 86.79% | 51.80% | 7 | 986.6 | 4835 | 4791 | JAFGTD01<sup>b</sup> | This study |
| Genus Ga0209509 | 3300009770_26 | Anaerobic biogas reactor | 54.58 ± 3.99 | 40.93 ± 8.14 | 2.50 Mbp | 2.70 Mbp | 90.94% | 65.20% | 1 | 1008 | 2291 | 2253 | Ga0123332<sup>a</sup> | 52 |
|  | 3300009773_26 |  |  |  | 2.56 Mbp | 2.75 Mbp | 90.73% | 65.20% | 2 | 1006 | 2346 | 2307 | Ga0123333<sup>a</sup> | 52 |
|  | 3300026194_8 |  |  |  | 3.76 Mbp | 3.88 Mbp | 91.48% | 66.60% | 2 | 1035 | 3370 | 3326 | Ga0209509<sup>a</sup> | 52 |
|  | 3300026195_8 |  |  |  | 3.83 Mbp | 3.95 Mbp | 91.56% | 66.50% | 5 | 1041 | 3418 | 3371 | Ga0209312<sup>a</sup> | 52 |
|  | 3300026198_13 |  |  |  | 2.52 Mbp | 2.82 Mbp | 91.52% | 65.70% | 1 | 997 | 2344 | 2310 | Ga0209313<sup>a</sup> | 52 |
|  | 3300026250_9 |  |  |  | 3.75 Mbp | 3.87 Mbp | 91.48% | 66.60% | 3 | 1040 | 3348 | 3300 | Ga0209612<sup>a</sup> | 52 |
|  | 3300026252_8 |  |  |  | 3.75 Mbp | 3.86 Mbp | 91.47% | 66.60% | 2 | 1041 | 3337 | 3291 | Ga0209722<sup>a</sup> | 52 |
|  | 3300026255_14 |  |  |  | 2.70 Mbp | 2.91 Mbp | 91.17% | 65.60% | 2 | 999 | 2502 | 2465 | Ga0209613<sup>a</sup> | 52 |
|  | 3300026290_9 |  |  |  | 3.86 Mbp | 3.98 Mbp | 91.35% | 66.20% | 3 | 1045 | 3425 | 3376 | Ga0209510<sup>a</sup> | 52 |
|  | 3300026311_36 |  |  |  | 2.69 Mbp | 2.78 Mbp | 90.30% | 64.90% | 3 | 1004 | 2466 | 2423 | Ga0209723<sup>a</sup> | 52 |
|  | Zgenome_919 | Zodlton Spring sediment |  |  | 4.75 Mbp | 4.98 Mbp | 91.97% | 66.50% | 0 | 996 | 3038 | 3001 | JAFGSS01<sup>b</sup> | This study |

a: Accession numbers correspond to Gold analysis project number (for MAGs from the recently released 52,515 genomes in the earth microbiome catalogue collection (ref) that were deposited in IMG/M database).
b: GenBank assembly accession number
\*: used with PI permission.

### Functional annotation

Protein-coding genes were annotated using Prodigal (v 2.50) (60). BlastKOALA (61) was used to assign KEGG orthologies (KO) to protein-coding genes, followed by metabolic pathways visualization in KEGG mapper (62). In addition, all genomes were queried with custom-built HMM profiles for sulfur metabolism, electron transport chain components (for alternate complex III), C1 metabolism, and hydrogenases. To construct hmm profiles, Uniprot reference sequences for genes with an assigned KO number were downloaded, aligned with Mafft (63), and the alignment was used to construct hmm profiles using the hmmbuild function of HMMer (v 3.1b2) (64). For genes without a designated KO number, a representative protein was queried against the KEGG genes database using Blastp, and hits with e-values <1e^−80^ were downloaded, aligned, and used to construct an hmm profile as described above. Hydrogenases hmm profiles were built using alignments downloaded from the Hydrogenase Database (HydDB) (65). The hmmscan function of HMMer (64) was used with the constructed profiles and a thresholding option of -T 100 to scan the protein-coding genes for possible hits. Further confirmation was achieved through phylogenetic assessment and tree building procedures. For that, putatively identified Acidobacteria sequences were aligned with Mafft (63) against the reference sequences used to build the HMM database, and the alignment was then used to construct a maximum-likelihood phylogenetic tree using FastTree (v 2.1.10) (66). Sequences that clustered with reference sequences were deemed to be true hits and were assigned a corresponding KO number or function. Carbohydrate active enzymes (CAZYmes) (including glycoside hydrolases [GHs], polysaccharide lyases [PLs], and carbohydrate esterases [CEs]) were identified by searching all ORFs from all genomes against the dbCAN hidden Markov models V9 (67, 68) (downloaded from the dbCAN web server in September 2020) using hmmscan. AntiSMASH 3.0 (69) was used with default parameters to predict biosynthetic gene clusters in the genomes. Canonical correspondence analysis (CCA) was used to identify the correlation between the genus, environmental source, and the types of BGCs predicted in the genomes using the function cca in the R package Vegan (https://cran.r-project.org/web/packages/vegan/index.html).

To evaluate the novelty of the NRPS and PKS clusters identified in family UBA6911 genomes, we queried the synthetic genes from these clusters against the NCBI nt database (Downloaded in April 2020). A threshold of 75% identity over 50% of the query length was used to determine the novelty of these genes.

### Phylogenetic analysis of XoxF methanol dehydrogenase and dissimilatory sulfite reductase DsrAB

Family UBA6911 predicted XoxF methanol dehydrogenase, as well as dissimilatory sulfite reductase subunits A and B were compared to reference sequences for phylogenetic placement. Family UBA6911 predicted XoxF protein sequences were aligned to corresponding reference sequences from other methylotrophic taxa, while the dissimilatory sulfite reductase subunits A, and B were aligned to corresponding subunits from sulfate-reducing taxa using Mafft (63). DsrA and DsrB alignments were concatenated in Mega X (70). The XoxF alignment, and the DsrAB concatenated alignment were used to construct maximum-likelihood phylogenetic trees using FastTree (v 2.1.10) (66).

### Ecological distribution

To further examine the ecological distribution of Family UBA6911 genera, we analyzed 177 near full length 16S rRNA gene sequences (>1200 bp) associated with this lineage in SILVA database (r138.1) (25) (Silva classification: Bacteria;Acidobacteriota;Subgroup 18;). A near-complete 16S rRNA gene from each genus (with the exception of genus QHZH01 represented by a single genomic assembly that unfortunately lacked a 16S rRNA gene) was selected as a representative and was included in the analysis. Sequences were aligned using the SINA aligner (71), and the alignment was used to construct maximum-likelihood phylogenetic trees with FastTree (66). The environmental source of hits clustering with the appropriate reference sequences were then classified with a scheme based on the GOLD ecosystem classification scheme (72). All phylogenetic trees were visualized and annotated in iTol (73).

### Sequence and MAG accessions

Metagenomic raw reads for Zodletone sediment are available under SRA accession SRX9813571. Zodletone whole genome shotgun project was submitted to GenBank under Bioproject ID PRJNA690107 and Biosample ID SAMN17269717. The individual assembled Acidobacteria MAGs analyzed in this study have been deposited at DDBJ/ENA/GenBank under the accession JAFGAO000000000, JAFGSS000000000, JAFGIY000000000, and JAFGTD000000000.

## Results

### Ecological distribution patterns of Family UBA6911

Family UBA6911 genomes clustered into 5 genera based on AAI and shared gene content values (Figure 1, Table 1). Genera Gp18_AA60 (n= 3) and QHZH01 (n=1) genomes were exclusively binned from a grassland meadow within the Angelo Coastal Range Reserve in Northern California (30). Genus Ga0209509 genomes were mostly (10/11 genomes) binned from an anaerobic gas digestor (WA, USA) (52). Genus Ga0212092 (n=2) genomes were binned from Cone pool hot spring microbial mat in California, USA. Finally, genus UBA6911 (10 genomes) displayed a broader distribution pattern as its genomes were recovered from multiple, mostly freshwater habitats, e.g. river and estuary sediments and Zodletone spring sediment (Table 1, Figure 1).

To further examine the global ecological distribution of Family UBA6911, we analyzed 177 near full-length 16S rRNA gene amplicons generated in multiple culture-independent amplicon-based diversity surveys and available through SILVA database (r138.1) (Figure 2). Genera distribution patterns gleaned from the origin of MAGs were generally confirmed: 16S rRNA sequences affiliated with genus Gp18_AA60 were mostly recovered from soil (14/15, 93.3%); those affiliated with the genus Ga0209509 were mostly encountered in anaerobic digestors (11/14, 85.7%); and the majority of 16S rRNA sequences affiliated with the genera Ga0212092 (90.9%) and UBA6911 (58.8%) were encountered in a wide range of freshwater environments (e.g. freshwater lake sediments, estuary sediments, and thermal springs). Collectively, the combined MAGs and 16S rRNA amplicon data suggest a preference for soil for genera Gp18_AA60 and QHZH01, a preference for anaerobic digestors for Ga0209509, and a wide occurrence of genera Ga0212092 and UBA6911 in freshwater habitats.

**Figure 2.**
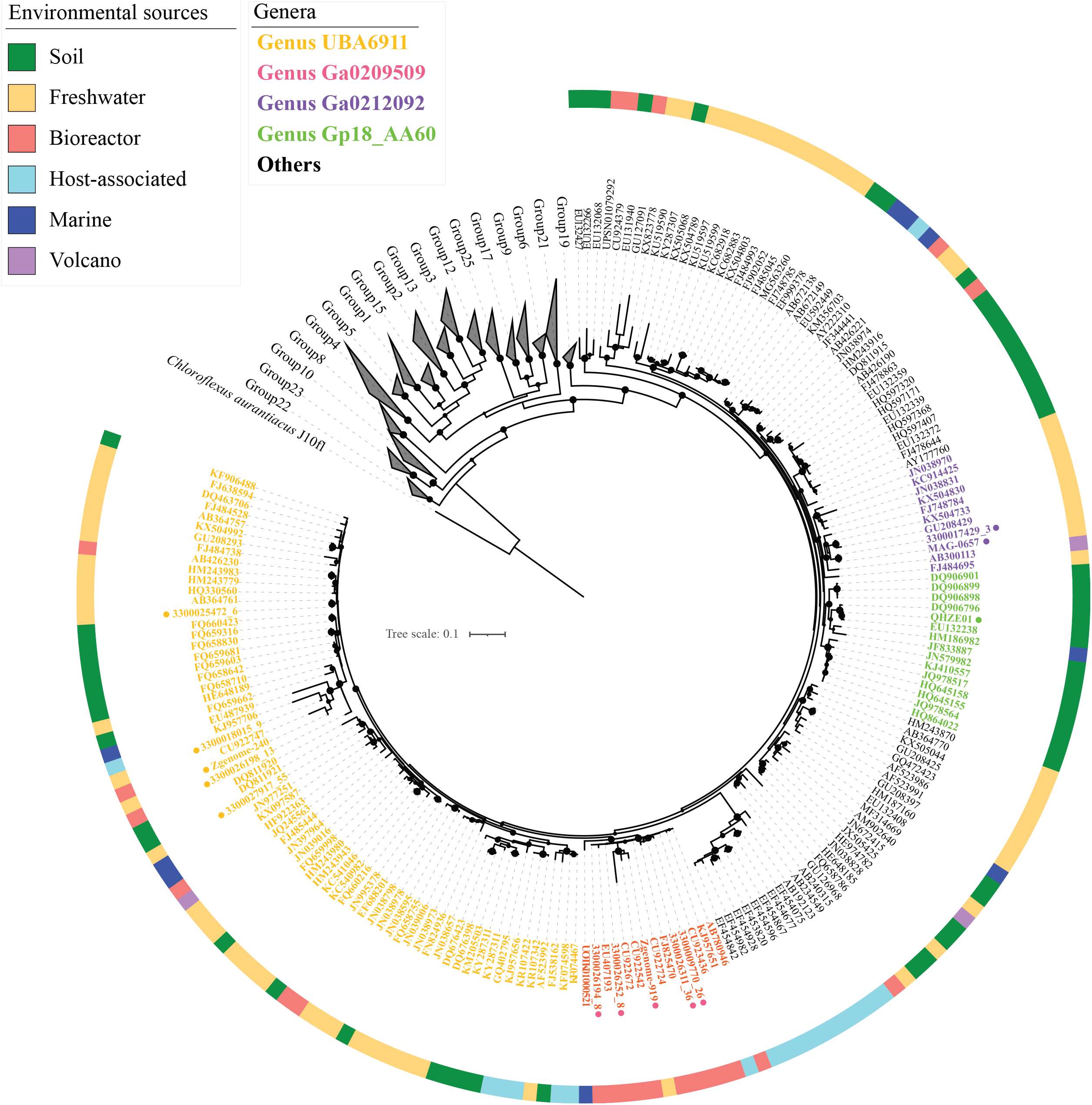
Phylogenetic tree based on 16S rRNA gene for Acidobacteria classes. Groups other than family UBA6911 (subgroup 18) are shown as grey wedges. A total of 177 near-full length (>1200 bp) 16S rRNA sequences belonging to Acidobacteria Group 18 in Silva database are shown with their GenBank accession numbers. Representative 16S rRNA sequences from the analyzed genomes are shown with their corresponding Bin name (as in Table 1), and with a colored dot next to the name for ease of recognition. Branch labels are color-coded by genus, as shown in the figure legend (using the same color scheme as in Figure 1). The track around the tree shows the environmental classification of the ecosystem from which the sequence was obtained and the corresponding color codes are shown in the figure legend. Sequences were aligned using the SINA aligner (71), and the alignment was used to construct maximum-likelihood phylogenetic tree with FastTree (66). Bootstrap values (from 100 bootstraps) are displayed as bubbles for branches with ≥70% support. The tree was rooted using *Chloroflexus aggregans* (GenBank accession number M34116.1) as the outgroup.

### General genomic and structural features of family UBA6911 genomes

Soil-affiliated genera (Gp18_AA60 and QHZH01) displayed larger estimated genome size (8.05 ± 0.32 Mbp), when compared to freshwater UBA6911, and Ga0212092 (5.45 ± 0.68 Mbp, and 4.45 ± 0.17 Mbp, respectively), and anaerobic digestor genus Ga0209509 (3.5 ± 0.75 Mbp) (Table 1). GC content was generally high in all genomes (Table 1), with three genera (QHZH01, Ga0209509, and Ga0212092) exhibiting >65% GC content. Structurally, all genera in family UBA6911 are predicted to be gram-negative rods with a similar predicted membrane phospholipid composition (Tables 2, and S3). Genes mediating type IV pilus formation, generally involved in a wide range of functions, e.g. adhesion and aggregation, twitching motility, DNA uptake and protein secretion (74), were observed in most genera. Secretion systems I and II were identified in all genomes. Interestingly, type VI secretion system, typically associated with pathogenic Gram-negative bacteria (mostly Proteobacteria), and known to mediate protein transport to adjacent cells as a mean of bacterial antagonism (75), was identified in all but a single (anaerobic digestor Ga0209509) genus. Searching the Annotree database (76) identified type VI secretion system genes in only 14 Acidobacteria genomes, all belonging to class Holophagae, suggesting the rare distribution of such trait in the phylum. Finally, genomes of the soil genus Gp18_AA60 possess an exceptionally large number of CRISPR genes (24.67 ± 6.5) (Table 1).

**Table 2.** Salient defining features of Family UBA6911 genera.

|  | Genus Gp18_AA60 | Genus QHZH01 | Genus UBA6911 | Genus Ga0209509 | Genus Ga0212092 |
| --- | --- | --- | --- | --- | --- |
| **Predicted structural features** |  |  |  |  |  |
| Flagellar motility | n | n | Y | n | Y |
| Type IV pilus assembly | Y | n | Y | Y | Y |
| Chemotaxis | n | n | Y | n | n |
| Type VI Secretion System | Y | Y | Y | n | Y |
| CRISPR count | 24.67 ± 6.5 | 2 | 5.4 ± 5.4 | 2.18 ± 1.33 | 4.5 ± 0.71 |
| **Biosynthesis: Biosynthetic gene clusters** |  |  |  |  |  |
| Terpenes | Y | Y | Y | Y | Y |
| Phenazine | Y | Y | n | n | n |
| NRPS/ PKS | Y | Y | Y | Y | Y |
| Homoserine lactones | n | n | n | Y | n |
| Bacteriocin | n | Y | Y | n | n |
| **Heterotrophic substrates predicted to support growth** |  |  |  |  |  |
| **Sugars<sup>a</sup>** |  |  |  |  |  |
| CAZymes (GHs + PLs + CEs) | 98.3 ± 11.7 | 166 | 48.2 ± 22.3 | 45.6 ± 13.4 | 42.5 ± 0.71 |
| Hexoses | Glu, Man, Gal, Fru | Glu, Man | Glu, Man, Gal, Fru | Glu, Man, Gal, Fru | Glu, Man, Gal |
| Hexosamines | Y | Y | Y | Y | Y |
| Hexuronic acids | n | n | Y | Y | n |
| Sialic acid | n | Y | n | n | n |
| Sugar alcohols | Ino, Sorb, Xylitol | Ino, Sorb, Xylitol | Sorb, Xylitol | Sorb, Xylitol | Xylitol |
| Pentoses | Lyx, Xyl | Lyx, Xyl | n | Xyl | n |
| **Amino acids** |  |  |  |  |  |
| Acidic and amides | D, N, E, Q | D, N, E, Q | D, N, E, Q | D, E, Q | D, N, E, Q |
| Aliphatic | A, G | A, G | A, G, V, L, I | A, G | A, G |
| Aromatic | n | n | Y | n | n |
| Basic | H | n | R, K | R | R, H |
| Hydroxy and S containing | S, T | S, T | M | M | S, T, M |
| Cyclic | P | P | P | P | P |
| **C1 compounds** |  |  |  |  |  |
| Methanol | n | Y | n | n | n |
| Methylamines | n | Y | n | n | n |
| Formaldehyde | n | Y | Y | n | n |
| **Predicted respiratory capacities** |  |  |  |  |  |
| Aerobic (low affinity) | Y | Y | Y | Y | Y |
| Aerobic (High affinity) | Y | n | Y | Y | Y |
| Dissimilatory nitrate reduction to ammonium | n | n | Y |  | Y |
| Dissimilatory nitrite reduction to ammonium | n | n |  | Y | n |
| Dissimilatory sulfate reduction | n | n | Y | n | Y |
| TMAO respiration | n | n | n | n | Y |
| **Predicted fermentation products** |  |  |  |  |  |
| Acetate | Y | Y | Y | Y | n |
| Ethanol | n | Y | Y | Y | Y |
| Formate | n | n | Y | Y | n |
| Acetone | n | n | Y | n | n |
| Acetoin and Butanediol | Y | n | Y | Y | Y |

a: Sugars are abbreviated as follows; Glu, glucose; Man, mannose; Gal, galactose; Fru, fructose; Ino, myo-inositol; Sorb, sorbitol; Lyx, lyxose; Xyl, xylose.
Y, full pathway identified; n, full pathway missing.

### Anabolic capabilities in family UBA6911 genomes

All UBA6911 genomes showed a fairly extensive anabolic repertoire, with capacity for biosynthesis of the majority of amino acids from precursors (from 15 in Genus Ga0212092 to 20 in genus UBA6911) (Table S3). Gluconeogenic capacity was observed in all genera (Table S3). Cofactors biosynthesis capability was also widespread in all but the freshwater Cone Pool Ga0212092 genomes (Table S3).

A complete assimilatory sulfate reduction pathway for sulfate uptake and assimilation was observed in all genomes. In addition, the presence of taurine dioxygenase (in soil genus QHZH01 genome) and alkanesulfonate monooxygenase (in soil genus gp18_AA60 genomes) argue for the capacity for organosulfur compounds assimilation. For nitrogen assimilation, in addition to NH_4_^+^ and amino acids, multiple pathways for N assimilation from organic substrates were identified. These include the presence of urease genes (*ureABC*) for urea assimilation in soil genus gp18_AA60 genomes. As well, genes for N extraction from arylformamides (arylformamidase, EC:3.5.1.9), N, N-dimethylformamide (N,N-dimethylformamidase EC:3.5.1.56), and N-formylglutamate (formylglutamate deformylase, EC:3.5.1.68) were identified to various degrees (Table S3) in all genera, with the exception of freshwater Cone Pool genus Ga0212092 genomes. No evidences for assimilatory nitrate reduction or nitrogen fixation were identified in any of the genomes.

In addition to the above biosynthetic capacities, all examined genomes, regardless of environmental source or genus-level affiliation, encoded biosynthetic gene clusters (BGC) ranging from 3 to 12. Such clusters include polyketide synthases (PKS) and non-ribosomal peptide synthases (NPRS), homoserine lactones, terpenes, phenazines, and bacteriocin. While the number of BGC per genome did not differ much between genera (Table S4), differences in the gene clusters, and hence putative products, were identified (Table 2, Figure S1). For example, phenazines biosynthetic clusters were only identified in the soil genera genomes, while BCGs for homoserine lactone biosynthesis were exclusive to the anaerobic digestor Ga0209509 genomes. While not exclusive, terpene biosynthetic clusters appear to be more enriched in anaerobic digestor Ga0209509 genomes, while NRPS and PKS clusters were enriched in the soil genera genomes and UBA6911 genomes. Finally, bacteriocin biosynthesis cluster was only identified in genus QHZH01 genome and a single genome affiliated with genus UBA6911 (Figure S1).

We queried NRPS and PKS clusters identified in all 28 genomes against the NCBI nt database. Surprisingly, only 14 genes (3.8%) had significant hits to previously deposited sequences (Table S5). Such high level of sequence novelty potentially implies the production of novel chemical compounds from such clusters.

### Substrate utilization patterns in family UBA6911 genomes

Genomic analysis of all Family UBA6911 genomes suggests a heterotrophic lifestyle with robust aminolytic and saccharolytic machineries. Genomes from all genera encoded an excellent capacity to metabolize proteins and amino acids. An arsenal of endopeptidaes, oligopeptidases, and dipeptidases were identified in all genomes (Table S6). Oligopeptide transporters (in UBA6911 genomes), peptide/Ni transporters (all genomes), and dedicated amino acid transporters (e.g. branched chain amino acid transporters in QHZH01, UBA6911, and Ga0212092 genomes, and trp/tyr transporters in gp18_AA60 genomes) suggest the capacity for amino acids and oligo/dipeptide uptake. Further, all genomes to various degrees showed degradation pathways for a wide range of amino acids, ranging from 9 (in genus Ga0212092) to 15 (in genus UBA6911) (Tables 2, and S3).

Similarly, with the notable exception of members of Freshwater Cone Pool Genus Ga0212092, a robust machinery for sugar metabolism was identified. (Tables 2, and S3). Glucose, mannose, galactose, fructose, hexosamines e.g., N-acetyl hexosamines, uronic acids, sorbitol, and xylitol degradation capacities were identified in all other four genera (Tables 2, and S3). Further, soil genera genomes encoded the full degradation machinery for *myo*-inositol (*iolBCDEG*), an abundant soil component (77). Interestingly, QHZH01 genome also encoded the full machinery for sialic acid (an integral component of soil fungal cell walls (78)) degradation to fructose-6-P.

Analysis of the carbohydrate active enzyme (CAZyme) repertoire was conducted to assess Family UBA6911 polysaccharide degradation capacities. A notable expansion of the CAZyme (CE + PL + GH) repertoire in soil genomes was observed (166 in QHZH01 genome, 98.3 ±11.7 in gp18_AA60 genomes), compared to 46.48 ± 17.01 in all other genera (Table S7). Higher numbers in soil genera genomes were brought about by the expansion of families GH33 (sialidase), GH165 (β-galactosidase), and GH56 (hyaluronidase) in genus QHZH01, of family GH13 (amylase) in gp18_AA, and of family GH109 (N-acetylhexosaminidase) in both genera (Table S7). As described above, genes encoding degradation machineries of the released monomer products for these enzymes (sialic acid, galactose, uronic acid, glucose, and N-acetylhexosamine) are encoded in the soil genomes (Tables 2, and S3).

Overall, all Family UBA6911 genomes analyzed in this study displayed a notable absence or extreme paucity of CAZy families encoding/initiating breakdown of large polymers, e.g. endo- and exo-cellulases (only 6 copies of GH5 genes, and 5 copies of GH9 genes in all 28 genomes examined), hemicellulases (only 19 copies of GH10 genes in all 28 genomes examined), and pectin degradation (only 22 copies of GH28 genes in all genomes examined), suggesting the limited ability of all members of the family for degrading these high molecular weight polysaccharides (Table S7). Collectively, the CAZyome and sugar degradation patterns of Family UBA6911 argue for a lineage specializing in sugar, but not polymer, degradation, with added specialized capacities for de-branching specific sugars from polymers in the soil-dwelling genera.

Further, methylotrophic C1 degradation capacity for methanol and methylamine utilization was observed in soil QHZH01 genome. The genome encoded a methanol dehydrogenase [EC:1.1.2.10] for methanol oxidation to formaldehyde (Figure 3, Tables 2, and S3). QHZH01 methanol dehydrogenase belonged to the lanthanide-dependent pyrroloquinoline quinone (PQQ) methanol dehydrogenase XoxF-type, and clustered closely with sequences from *Candidatus* Methylomirabilis (NC10) and Verrucomicrobia (Figure 4). The accessory *xoxG* (c-type cytochrome) and *xoxJ* (periplasmic binding) genes were fused and encoded in QHZH01 genome downstream of the *xoxF* gene. A quinohemoprotein amine dehydrogenase [EC:1.4.9.1] for methylamine oxidation to formaldehyde (Figure 3, 4, Tables 2, and S3) was also identified in the genome. Further, the genome also encoded subsequent formaldehyde oxidation to formate via the glutathione-independent formaldehyde dehydrogenase [EC:1.2.1.46], as well as formate oxidation to CO_2_ via formate dehydrogenase [EC:1.17.1.9]. Finally, for assimilating formaldehyde into biomass, genes encoding the majority of enzymes of the serine cycle were identified the genome, as well as the majority of genes encoding the ethylmalonyl-CoA pathway for glyoxylate regeneration (Figure 3, Tables 2, and S3).

**Figure 3.**
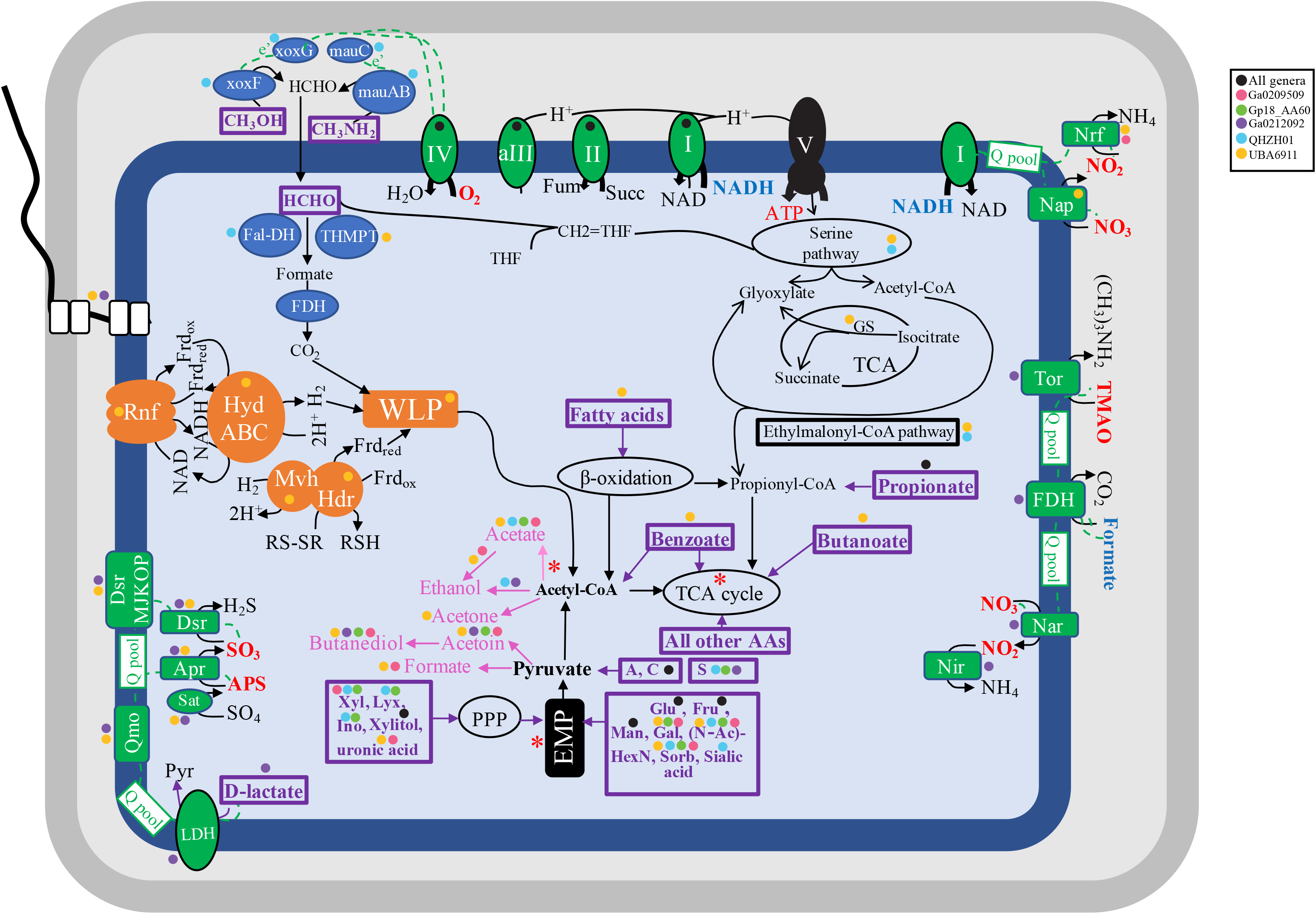
Cartoon depicting different metabolic capabilities encoded in Family UBA6911 genomes with capabilities predicted for different genera shown as colored circles (all orders, black; Genus Ga0209509, pink; Genus Gp18_AA60, green; Genus Ga0212092, purple; Genus QHZH01, cyan; Genus UBA6911, yellow). Enzymes for C1 metabolism are shown in blue. Electron transport components are shown in green and electron transfer is shown as dotted green lines from electron donors (shown in boldface blue text) to terminal electron acceptors (shown in boldface red text). The sites of proton extrusion to the periplasm are shown as black arrows, as is the F-type ATP synthase (V). Proton motive force generation, as well as electron carrier recycling pathways associated with the operation of the WLP are shown in orange. All substrates predicted to support growth are shown in boldface purple text within thick purple boxes. Fermentation end products are shown in pink. Sites of substrate level phosphorylation are shown as red asterisks. A flagellum is depicted, the biosynthetic genes of which were identified in genomes belonging to the genera UBA6911 and Ga0212092. The cell is depicted as rod-shaped based on the identification of the rod shape determining proteins *rodA, mreB* and *mreC* in all genomes. Abbreviations: Apr, the enzyme complex adenylylsulfate reductase [EC:1.8.99.2]; APS, adenylyl sulfate; Dsr, dissimilatory sulfite reductase [EC:1.8.99.5]; EMP, Embden Meyerhoff Paranas pathway; Fal-DH, formaldehyde dehydrogenase; FDH, formate dehydrogenase; Frd_ox/red_, Ferredoxin (oxidized/ reduced); Fru, fructose; fum, fumarate; Gal, galactose; Glu, glucose; GS, glyoxylate shunt; Hdr, heterodisulfide reductase complex; HydABC, cytoplasmic [Fe Fe] hydrogenase; I, II, aIII, and IV, aerobic respiratory chain comprising complexes I, II, alternate complex III, and complex IV; Ino, *myo*-inositol; LDH, L-lactate dehydrogenase; Lyx, lyxose; Man, mannose; mauABC; methylamine dehydrogenase; Mvh, Cytoplasmic [Ni Fe] hydrogenase; Nap, nitrate reductase (cytochrome); Nar, nitrate reductase; Nir, nitrite reductase (NADH); Nrf, nitrite reductase (cytochrome c-552); N-AcHexN, N-acetylhexosamines; PPP, pentose phosphate pathway; Pyr, pyruvate; Q pool, quinone pool; Qmo, quinone-interacting membrane-bound oxidoreductase complex; RNF, membrane-bound RNF complex; RSH/RS-SR, reduced/oxidized disulfide; Sorb, sorbitol; succ, succinate; TCA, tricarboxylic acid cycle; THMPT, tetrahydromethanopterin-linked formaldehyde dehydrogenase; TMAO, trimethylamine N-oxide; Tor, trimethylamine-N-oxide reductase (cytochrome c); V, ATP synthase complex; WLP, Wood Ljungdahl pathway; xoxFG, methanol dehydrogenase; Xyl, xylose.

**Figure 4.**
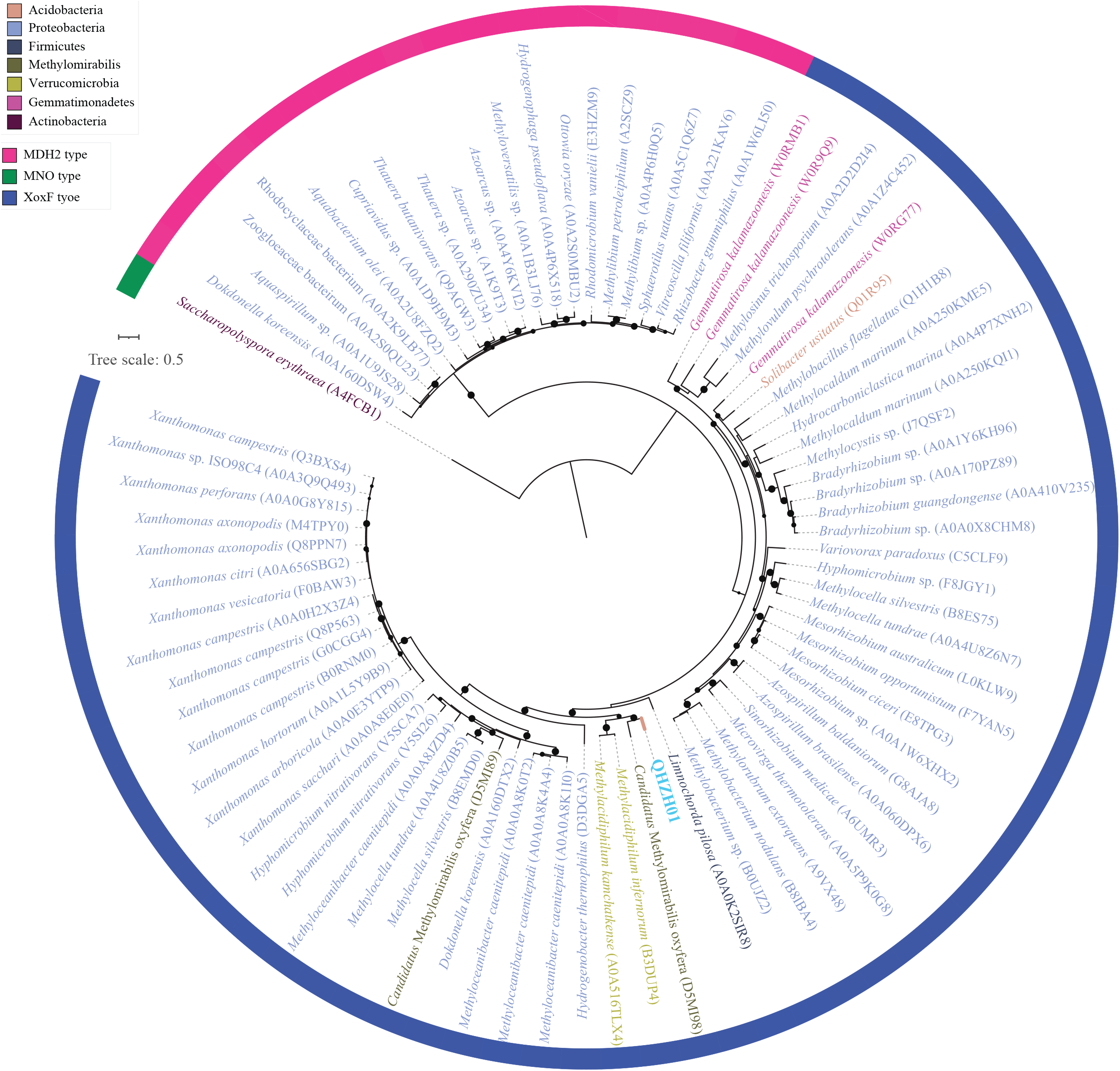
Phylogenetic affiliation for Family UBA6911 methanol dehydrogenase (XoxF) in relation to reference sequences. Genus QHZH01 sequence is shown in cyan. Uniprot accessions are shown for reference sequences that are color coded by phylum as shown in the legend. Bootstrap support values are based on 100 replicates and are shown for nodes with >70% support. The track around the tree corresponds to the family of methanol dehydrogenase and is color coded as shown in the figure legend.

In addition, several genomes in the genus UBA6911 encoded formaldehyde oxidation (via tetrahydromethopterin-linked pathway, glutathione-independent formaldehyde dehydrogenase [EC:1.2.1.46], and glutathione-dependent formaldehyde dehydrogenase [EC:1.1.1.284, EC 3.1.2.12]) to formate, formate oxidation to CO_2_ via formate dehydrogenase [EC:1.17.1.9], formaldehyde assimilation into biomass (genes encoding the majority of enzymes of the serine cycle were identified in some genomes), as well as glyoxylate regeneration (the majority of genes encoding the ethylmalonyl-CoA pathway were identified in some genomes, with only one genome encoding glyoxylate shunt genes) (Figure 3, Tables 2, and S3). Surprisingly, upstream genes for formaldehyde production from C1 compounds (e.g. methane, methanol, methylamine, C1 sulfur compounds) were missing from all genomes.

Finally, in addition to proteins, carbohydrates, and C1 compounds, other potential carbon sources for Family UBA6911 were identified. These include long chain fatty acids (a complete beta oxidation pathway) in genus UBA6911 genomes, short chain aliphatic fatty acids (propionate and butyrate degradation to acetyl-CoA) in both UBA6911 and Ga0212092 genomes, and benzoate in genus UBA6911 genomes (Table S3).

### Respiratory capacities

All Family UBA6911 genomes encoded respiratory capacities. Genomic analysis suggested that the soil genera Gp18_AA60 and QHZH01 utilize O_2_ as their sole electron acceptor, based on their possession of a respiratory chain comprising complex I, II, alternate complex III, and IV (cytochrome oxidase aa3) (Tables 2 and S3). On the other hand, all anaerobic digestor genus Ga0209509 genomes could mediate dissimilatory nitrite reduction to ammonium, based on the presence of nitrite reductase (cytochrome c-552) [EC:1.7.2.2], plus respiratory complexes I and II. Only 6 out of the 11 genomes in this genus possessed a complete aerobic respiratory chain. Freshwater genus Ga0212092 genomes from Cone pool microbial sediments were notable in encoding a complete aerobic respiratory chain as well as the complete machinery for dissimilatory sulfate reduction. Additional plausible anaerobic electron acceptor for genus Ga0212092 is trimethylamine N-oxide (TMAO). Here, electron transfer is thought to occur from formate via the membrane-bound formate dehydrogenase [EC: 1.17.1.9] through the quinone pool onto trimethylamine N-oxide (TMAO) reductase [EC: 1.7.2.3] eventually reducing trimethylamine N-oxide to trimethylamine (Figure 3, Tables 2, and S3), as previously shown in *E. coli* (79) and *Rhodopseudomonas capsulata* (80). Finally, members of the freshwater genus UBA6911 demonstrated the most versatile respiratory capacities, with evidence for aerobic respiration (in all genomes except a single Oak River estuary sediment MAG), dissimilatory nitrite reduction to ammonium in genomes from Washington anaerobic gas digestor and Zodletone spring sediment, as well as dissimilatory sulfate reduction to sulfide in Noosa River sediment, and Zodletone spring sediment genomes (Figure 3, Tables 2, and S3).

Sulfate-reduction machinery identified in the genomes of freshwater genera UBA6911, and Ga0212092 included 3’-phosphoadenosine 5’-phosphosulfate synthase [Sat; EC:2.7.7.4 2.7.1.25] for sulfate activation to adenylyl sulfate (APS), the enzyme complex adenylylsulfate reductase [AprAB; EC:1.8.99.2] for APS reduction to sulfite, the quinone-interacting membrane-bound oxidoreductase complex [QmoABC] for electron transfer, the enzyme dissimilatory sulfite reductase [DsrAB; EC:1.8.99.5] and its co-substrate DsrC for dissimilatory sulfite reduction to sulfide, and the sulfite reduction-associated membrane complex DsrMKJOP for linking cytoplasmic sulfite reduction to energy conservation (Figure 3). Phylogenetic affiliation using a concatenated alignment of DsrA and DsrB proteins placed genus UBA6911 sequences close to Thermoanaerobaculia Acidobacteria sequences from hydrothermal vents (Figure 5), while genus Ga0212092 sequences were close to Group 3 Acidobacteria sequences from hydrothermal vents and peatland soil (Figure 5).

**Figure 5.**
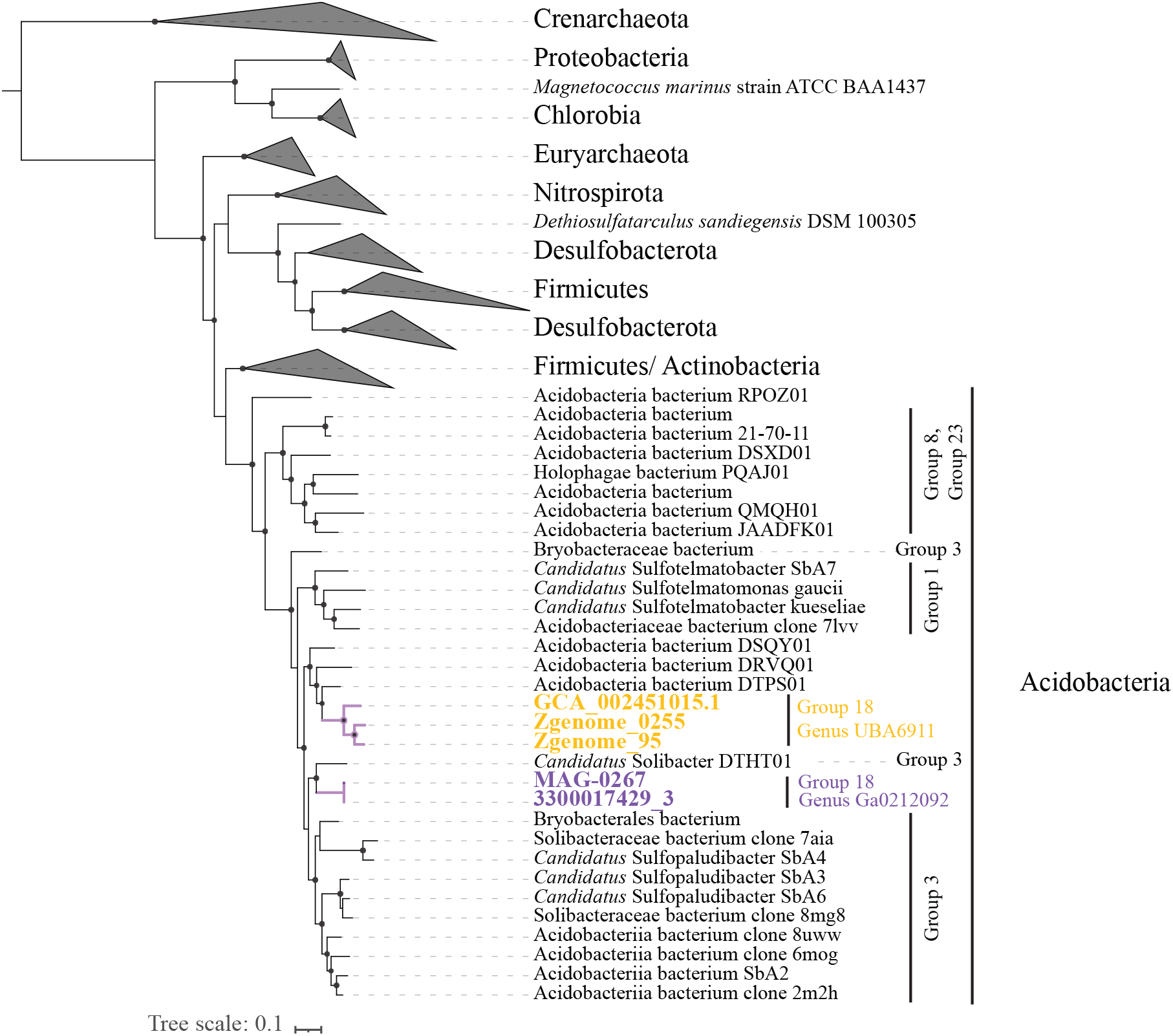
Maximum likelihood phylogenetic tree based on the concatenated alignment of the alpha and beta subunits of dissimilatory sulfite reductase (DsrAB) from Family UBA6911 in relation to reference sequences. Genus UBA6911 sequences are shown in yellow, while genus Ga0212092 sequences are shown in purple. Reference sequences from phyla other than Acidobacteria are shown as grey wedges. Acidobacteria references are labeled by the subgroup number. Bootstrap support values are based on 100 replicates and are shown for nodes with >70% support.

Finally, in addition to inorganic electron acceptors, the majority of genus UBA6911 genomes encoded a complete Wood-Ljungdahl (WL) pathway (Figure 3). WL is a versatile widespread pathway that is incorporated in the metabolic schemes of a wide range of phylogenetically disparate anaerobic prokaryotes e.g. homoacetogenic bacteria, hydrogenotrophic methanogens, autotrophic sulfate-reducing prokaryotes, heterotrophic sulfate-reducing bacteria, syntrophic acetate-oxidizing (SAO) bacteria, as well as acetoclastic methanogens. When operating in the reductive direction, the pathway can be used for carbon dioxide fixation and energy conservation during autotrophic growth, or as an electron sink during heterotrophic fermentative metabolism. When operating in the oxidative direction, the pathway is used for acetate catabolism (81–83). Syntrophic acetate oxidizers employing the oxidative WLP usually possess high affinity acetate transporters to allow for the uptake of small concentrations of acetate, a competitive advantage in the presence of acetoclastic methanogens (84). The absence of genes encoding high affinity acetate transporters in any of the genomes argues against the involvement of WLP in syntrophic acetate catabolism. The possibility of its operation for autotrophic CO_2_ fixation is also unlikely due to the absence of evidence for utilization of an inorganic electron donor (e.g. molecular H_2_). Its most plausible function, therefore, is acting as an electron sink to re-oxidize reduced ferredoxin, as previously noted in *Candidatus* Bipolaricaulota genomes (84). The RNF complex encoded in the majority of genus UBA6911 genomes would allow the re-oxidation of reduced ferredoxin at the expense of NAD, with the concomitant export of protons to the periplasm, thus achieving redox balance between heterotrophic substrate oxidation and the WL function as the electron sink. Additional ATP production via oxidative phosphorylation following the generation of the proton-motive force is expected to occur via the F-type ATP synthase encoded in all genomes. Recycling of electron carriers would further be achieved by the cytoplasmic electron bifurcating mechanism HydABC plus MvhAGD-HdrABC, both of which are encoded in the genomes (Figure 3).

### Fermentative capacities

In addition to respiration, elements of fermentation were also encoded in all genomes. All MAGs encoded pyruvate dehydrogenase, as well as 2-oxoacid ferredoxin oxidoreductase for pyruvate oxidation to acetyl-CoA. Genes encoding fermentative enzymes included the acetate production genes (acetyl-CoA synthase [EC:6.2.1.1] in the soil genera and UBA6911 genomes, acetate:CoA ligase (ADP-forming) [EC:6.2.1.13] in UBA6911 and Ga0209509 genera genomes, and phosphate acetyltransferase [EC:2.3.1.8] and acetate kinase [EC:2.7.2.1] (this latter pathway is associated with substrate level phosphorylation and was only encoded in UBA6911 genomes)), genes for ethanol production from acetate (aldehyde dehydrogenase [EC:1.2.1.3], and alcohol dehydrogenases) in UBA6911 and Ga0209509 genera genomes, genes for ethanol production from acetyl-CoA (aldehyde dehydrogenase [EC:1.2.1.10], and alcohol dehydrogenases) in QHZH01 and Ga0212092 genomes, genes for formate production from pyruvate (formate C-acetyltransferase [EC:2.3.1.54] and its activating enzyme) in UBA6911 and Ga0209509 genera genomes, genes for acetoin and butanediol production from pyruvate in all but QHZH01 genomes, and genes for acetone production from acetyl-CoA (acetyl-CoA C-acetyltransferase [EC:2.3.1.9], acetate CoA/acetoacetate CoA-transferase alpha subunit [EC:2.8.3.8 2.8.3.9], and acetoacetate decarboxylase [EC:4.1.1.4]) in UBA6911 genomes (Figure 3, Tables 2, and S3).

## Discussion

Our analysis reveals multiple defining features in all examined family UBA9611 genomes regardless of their origin or genus-level affiliation. All members of the family are predicted to be heterotrophs, with a robust anabolic capacity as well as a high level of catabolic versatility. Catabolism of a wide range of sugars, proteins, and amino acids by members of this family is predicted. As well, all members of this family appear to possess respiratory capacities. Such traits are similar to known metabolic capacities of the majority of examined genomes within the Acidobacteria (15, 33, 85). However, our analysis uncovered multiple interesting differences between the examined genomes, all of which appear to be genus-specific and, given the strong preference of various genera to specific habitats, habitat-specific. Such differences appear to differentiate the two soil genera from the three non-soil (freshwater and anaerobic digestor) genera (Tables 2, S3, Figure 3), and hence provide clues to the genus-level adaptation to the terrestrialization process within the family UBA6911. Three main differences are notable: Genomic architecture, substrate level preferences, and respiratory/electron accepting processes.

Soil genera gp18_AA60 and QHZH01 possess significantly larger genome size (8.05 ± 0.32 Mbp), compared to freshwater genomes (5.28 ± 0.73 Mbp), and anaerobic digestor genomes (3.5 ± 0.75 Mbp) (Table 1). Soil Acidobacteria usually (33, 85, 86), but not always (33, 87, 88) exhibit large genome sizes, compared to non-soil Acidobacteria (e.g. (35, 36) for a review see (89)) strongly arguing that genome expansion is associated with terrestrialization in the Acidobacteria. The association between genome size expansion and adaptation to soils has been identified as a general trait across the microbial world (90), including Acidobacteria (85, 89). Such expansion has been attributed to a higher number of paralogous genes as a mechanism for new functions generation, or optimal resource utilization under the different environmental conditions occurring in the highly complex and spatiotemporally dynamic soil ecosystem (91). Our analysis suggests that such expansion in gp18_AA60 and QHZH01 genomes is due to a broad increase in the number of genes in soil genera across a wide range of cellular/metabolic functions, rather than gene duplication or differences in coding density. In addition to genome size expansion, soil genus gp18_AA60 exhibited a high number of CRISPR loci (24.67 ± 6.5) (Table 1). In bacteria, CRISPR repeats arise from invading genetic elements that are incorporated into the host’s CRISPR locus. These short sequence tags (spacer sequences) are subsequently transcribed into small RNAs to guide the destruction of foreign genetic material (92). Within the Acidobacteria, examining 177 genomes (including MAGs and SAGs) affiliated with the phylum Acidobacteria in the IMG database revealed that 67 possessed at least 1 CRISPR count, and only four of these possessed >5 CRISPRs. The expansion in the number of CRISPR loci in gp18_AA60 genomes is possibly a protective mechanism against higher potential for viral infection by Acidobacteria-specific viruses in soil, which is expected given the higher relative abundance of members of this phylum in soil (23) compared to their low abundance in other habitats.

Multiple differences in catabolic capacities between soil and non-soil family UBA6911 genera were identified in this study. First, a larger CAZyome was identified in the soil genera, gp18_AA60 and QHZH01. This was mainly driven by the expansion of specific GH families, e.g. GH33 (sialidase), GH165 (β-galactosidase), and GH56 (hyaluronidase) in genus QHZH01, GH13 (amylase) in gp18_AA60, and GH109 (N-acetylhexosaminidase) in both genera. It is notable that the capacity for degradation of all released monomers (sialic acid, galactose, uronic acids, glucose, and N-acetylhexosamines) are indeed encoded in these soil genomes. As such acquisition of these specific CAZymes appears to be important for niche adaptation in family UBA6911 soil genomes. Second, the capacity for methanol and methylamine metabolism was predicted in members of the soil genus QHZH01. As previously noted (93), methylotrophy requires the possession of three metabolic modules for: C1 oxidation to formaldehyde, formaldehyde oxidation to CO_2_, and formaldehyde assimilation. QHZH01 encodes genes for all three modules. Formaldehyde assimilation via the serine cycle requires regeneration of glyoxylate from acetyl-CoA to restore glycine and close the cycle. In addition to all three modules described above, the QHZH01 genome also encodes the ethylmalonyl-CoA pathway for glyoxylate regeneration. Soil represent a major source of global methanol emissions (94), where demethylation reactions associated with pectin and other plant polysaccharides degradation contribute to the soil methanol pool. QHZH01 methanol dehydrogenase belongs to the xoxF family (Figure 4), previously detected in 187 genomes, including Acidobacteria SD1, SD5, and SD6 genomes (30), binned from Angelo soil, and identified as one of the most abundant proteins in a proteomics study from the same site (95). In contrast, some genus UBA6911 genomes possessed capacities for formaldehyde degradation and assimilation, but lacked any genes or modules for conversion of other C1 substrates to formaldehyde. The functionality and value of such truncated C1 machinery remains unclear. Formaldehyde in freshwater environments could be present as a contaminant in aquaculture facilities effluent (96), or as a product of C1 metabolism by other members of the community.

Finally, while all family UBA6911 genomes possessed respiratory chains, distinct differences in predicted respiratory capacities were observed. Soil genera gp18_AA60 and QHZH01 only encoded an aerobic respiratory ETC. In contrast, all anaerobic digestor genus Ga0209509 genomes encoded genes for dissimilatory nitrite reduction to ammonium, suggesting a capacity and/or preference to grow in strict anaerobic settings. Freshwater genera Ga0212092 and UBA6911 were the most diverse, combining aerobic capacity (in all but 1 genome), dissimilatory nitrite reduction to ammonium (in 4 UBA6911 genomes), trimethylamine N-oxide respiration (in both Cone Pool Ga0212092 genomes), potential use of WLP for electron acceptance (in genus UBA6911 genomes), and dissimilarity sulfate reduction capacities (in both Cone Pool Ga0212092 and some of UBA6911 genomes). This versatility could be a reflection of the relatively broader habitats where these genomes were binned, many of which exhibiting seasonal and diel fluctuation in O_2_ and other electron acceptors levels. Of note, is the presence of the complete machinery for dissimilatory sulfate-reduction, a process long thought of as a specialty for Deltaproteobacteria (Now Desulfobacterota (97)), some Firmicutes, and few thermophilic bacteria and archaea (98). However, with the recent accumulation of metagenomics data, genes for dissimilatory sulfate-reduction were identified in a wide range of phyla (2, 97). In the Acidobacteria, the presence and activity of genes for sulfate reduction has been reported from peatland samples (Subdivisions 1, and 3) (28). Subsequently, dissimilarity sulfate reduction capacities were identified in more Acidobacteria MAGs (Subdivision 23) from mine drainage (2) and recently, from marine fjord sediments of Svalbard (38). Our study adds Acidobacteria family UBA6911 (subgroup 18) to the list of the DSR-harboring Acidobacteria and suggest a role in freshwater habitats. The concurrent occurrence of dissimilatory sulfate reduction and aerobic respiration in the same MAG is notable, but has been previously reported in the Subdivision 23 Acidobacteria MAGs from marine fjord sediments of Svalbard (38). All Acidobacteria DsrAB sequences (Subdivisions 1, 3, 18, and 23) cluster within the reductive bacterial-type branches in the DsrAB concatenated phylogenetic trees, away from the oxidative Chlorobia and Proteobacteria (Figure 5). Also, the absence of DsrEFH homologs, known to be involved in the reverse DSR pathway, argue for the involvement of Acidobacteria DsrAB in the reductive dissimilatory sulfate reduction.

## Acknowledgments

This work has been supported by NSF grant 2016423 to NHY and MSE.

